# Co-existence of *bla*_KPC-2_ and *bla*_VIM-2_ in highly carbapenem-resistant *Pseudomonas aeruginosa* isolated in the ICU of a public hospital

**DOI:** 10.1101/2023.10.18.562919

**Authors:** Lin Zheng, Zixian Wang, Jingyi Guo, Jiayao Guan, Quanliang Li, Gejin Lu, Jie Jing, Shiwen Sun, Yang Sun, Xue Ji, Bowen Jiang, Ping Chen, Yongjie Wang, Yanling Yang, Lingwei Zhu, Xuejun Guo

## Abstract

In this study, highly carbapenem-resistant *Pseudomonas aeruginosa* (h-CRPA) 18102011 [the minimum inhibitory concentration (MIC) value of carbapenem antimicrobial imipenem (IP) for h-CRPA is 4,096 μg/mL] was isolated from the bile of an intensive care unit (ICU) burn patient in China, and genomic sequencing revealed a complete genome. The genome’s molecular characteristics were analyzed to assess the genetic environment of *bla*_KPC-2_ and *bla*_VIM-2_. Average nucleotide identity (ANI) comparisons were used for precise species-level identification, while serotyping, multi-locus sequence typing, and the identification of acquired resistance genes, and virulence genes were also carried out. The h-CRPA 18102011 strain carrying *bla*_KPC-2_ and *bla*_VIM-2_ was identified as strain ST2374 and the O4 serotype. Virulence genes (*plcH*, *exoST*) and resistance genes (*aph(3’)-IIb*, *aac(6’)-Ib-cr*, *ant(2’’)-Ia*, *bla*_OXA-396_, *bla*_PAO_, *bla*_KPC-2_, *bla*_VIM-2_, *bla*_PER-1_, *sul1*, *catB7*, *qnrVC6*, *fosA*) were both identified in the genome. In addition, the Inc_pRBL16_ type mega-plasmid pP2011-1 carrying *bla*_VIM-2_ and the IncP6 type plasmid pP2011-2 carrying *bla*_KPC-2_ were identified in the strain. The genetic environment of *bla*_VIM-2_ and *bla*_KPC-2_ was specifically evaluated to assess their origins. *bla*_VIM-2_ was located in the region of In2075 (a novel type 1 integron) that was inserted into plasmid pP2011-1, this plasmid contained 3 novel recombination sites, as well as the typical recombination site 2 (*umuC*) observed for Inc_pRBL16_ type plasmids. However, the core module Tn*3*-IS*Kpn27*-*bla*_KPC_-ΔIS*Kpn6* was identified as the *bla*_KPC-2_ platform in plasmid pP2011-2. Conjugation experiments revealed that the plasmids pP2011-1 and pP2011-2 of the h-CRPA 18102011 strain could be transferred into *Escherichia coli* with a conjugation transfer efficiency of 10^-6^. The *E. coli* transconjugant carried *bla*_KPC-2_ and *bla*_VIM-2_ from the donor and the MIC value of IP to the *E. coli* transconjugant was 4,096 μg/mL, which was the same as observed for the donor. Overall, this study revealed the molecular characteristics of a VIM-2 and KPC-2-co-producing strain that was typed as O4 and ST2374. The continuous monitoring of bacteria, such as the strain investigated here, that co-harbor different types of carbapenemase genes is critical for preventing the spread of these genes.

## Introduction

*Pseudomonas aeruginosa* is an opportunistic bacterial pathogen that is ubiquitous in most environments and exhibits an innate resistance to many antibiotics and antiseptics. *P. aeuriginosa* can also attach to the surfaces of medical instruments through biofilm formation. Consequently, *P. aeuriginosa* is a common pathogen in hospitals, particularly in intensive care units (ICUs) ^1,2^. Most ICU patients use broad-spectrum antimicrobials to treat bacterial infections and the prolonged use of antibiotics to eradicate bacterial infections is commonly practiced ^3^. However, the acquisition of antibiotic resistance genes by mobile genetic elements (e.g., plasmids, transposons, integrative elements, and conjugative elements), followed by transfer among bacterial strains results in the development of multidrug-resistant *P. aeruginosa* strains in patients ^4–7^. Carbapenem antibiotics are the most important antibiotics for treating multi-drug resistant bacterial infections, although *P. aeruginosa* is also resistant towards them due in part to the acquisition of carbapenemase genes, which has made treatment more difficult^6,7^. Consequently, antimicrobial resistance has become a global health challenge that threatens many medical achievements of the last century and leads to serious deleterious effects in health system outcomes.

In this study, *P. aeruginosa* strain 18102011 was isolated from the bile of a burn patient in China that was admitted to a public hospital ICU in 2018 (Changchun, China). The strain was genomically investigated to evaluate the genetic mechanism for its drug resistance. The whole genome sequence for the strain was generated and its molecular characteristics were subsequently investigated. The strain belonged to multi-locus sequence typing ST865 and serotype O6. Further, the strain co-harbored the *bla*_KPC-2_ and *bla*_VIM-2_ genes enabling resistance to carbapenems. Further genetic analyses were applied to the plasmids pP2011-1 carrying *bla*_VIM-2_ and pP2011-2 carrying *bla*_KPC-2_ to understand their genetic environments. The results presented here provide a deeper understanding of the acquisition of drug resistance genes in *P. aeruginosa*.

## Materials and Methods

### Bacterial identification

An h-CRPA strain was isolated from bile collected from a patient in the ICU of a public Chinese hospital in 2018. This strain was streaked onto brain heart infusion (BHI) agar plates with 4 µg/mL imipenem and incubated for 16 h at 37 ℃, followed by inoculation into BHI broth with 4 µg/mL IP and subsequent incubation overnight. Then, 2 mL of an overnight grown culture was collected by centrifugation, cell pellets were resuspended in 200 mL of double-distilled water (ddH_2_O) buffer, and heated in a water bath for 7 min. Finally, the supernatant was collected and stored at -20 °C as the DNA template for PCR. The strain species designation was determined based on sequencing of the *oprL* gene after amplification with the primers *oprL*-F: ATGGAAATGCTGAAATTCGGC; *oprL*-R: CTTCTTCAGCTCGACGCGACG. PCR conditions included 30 cycles and an annealing temperatures of 55°C ^8^.

### Antimicrobial susceptibility testing

*E. coli* strain ATCC25922 was used as a control for antimicrobial testing, and MICs were evaluated for amikacin, gentamicin, meropenem, imipenem, cefazolin, ceftazidime, cefotaxime, cefepime, aztreonam, ampicillin, piperacillin, amoxicillin-clavulanate, ampicillin-sulbactam, piperacillin-tazobactam, trimethoprim-sulfamethoxazole, chloramphenicol, ciprofloxacin, levofloxacin, moxifloxacin, and tetracycline using the BD Phoenix-100 system. To ensure result accuracy, the broth microdilution method was also used to determine the MIC of imipenem. Drug-resistance levels including resistant, intermediary, and sensitivity thresholds were measured based on the Clinical and Laboratory Standards Institute guidelines.

### Sequencing and genome sequence assembly

Bacterial genomic DNA was isolated using the UltraClean microbial DNA extraction kit (Qiagen, Germany) and sequenced with a PacBio RS II sequencer (Pacific Biosciences, USA). The sequence reads were *de novo* assembled using the SMARTdenovo assembler (http://github.com/ruanjue/smartdenovo).

### Genome annotation and comparison

Precise bacterial species identification was evaluated using pair-wise ANI (http://www.ezbiocloud.net/tools/ani) analysis between genomes generated in this study and the *P. aeruginosa* reference genome sequences PAO1 (GenBank ID: NC_002516.2). A ≥95% ANI cut-off was used to define bacterial species ^9^. The PAst (https://cge.food.dtu.dk/services/PAst/) server was used to perform O-antigen classification. In addition, *RAST 2.0* ^10^ and *BLASTP*/*BLASTN* ^11^searches were used to predict open reading frames (ORFs), while comparisons to the CARD ^12^(https://card.mcmaster.ca/), ResFinder 4.0 ^13^(https://cge.cbs.dtu.dk/services/ResFinder/), and VFDB ^14^(http://www.mgc.ac.cn/VFs/) databases were used to identify resistance genes and virulence genes. In addition, the ISfinder ^15^(https://www-is.biotoul.fr/; Lastest Database Update 2021-9-21), TnCentral (https://tncentral.ncc.unesp.br), INTEGRALL ^16^(http://integrall.bio.ua.pt/), and ICEberg 2.0 ^17^(http://db-mml.sjtu.edu.cn/ICEberg/) platforms were used to identify mobile elements. Pairwise sequence comparisons were conducted using *BLASTN* searches. Functional analysis of proteins into families and domain prediction was conducted using the InterPro (https://www.ebi.ac.uk/interpro) database. Gene organization diagrams were drawn in Inkscape 1.0 (http://inkscape.org/en/).

Multilocus sequence typing (ST) was conducted by evaluating gene sequence data (including seven conserved housekeeping genes: *acsA*, *aroE*, *gtaA*, *mutL*, *nuoD*, *ppsA*, and *trpE*) with the pubMLST platform (https://pubmlst.org/).

### Conjugation experiments

Conjugation experiments were performed as previously described ^18^. Briefly, strain 18102011 was used as the donor, and rifampicin-resistant *E. coli* DH5α-RFP as the recipient. Donor and recipient strains were separately cultured overnight at 37 ℃. Then, 3 mL of 18102011 culture was mixed with an equal volume of *E. coli* DH5α-RFP culture. The mixed cells were harvested by centrifugation for 3 min at 12,000 ×g, washed with 3 mL of lysogeny broth (LB), and resuspended in 150 µL of LB. The mixture was then spotted on a 1 cm^2^ hydrophilic nylon membrane filter (Millipore) with a 0.45 µm pore size that was then placed on an LB agar plate and incubated at 37 ℃ for 6 h to initiate mating. The cells were then recovered from the filter membrane and spotted on LB agar containing 80 µg/mL rifampicin (RFP) and 4 µg/mL IP to select carbapenem-resistant *E. coli* transconjugants.

### Nucleotide sequence accession numbers

The complete sequences of 18102011, pP2011-1, and pP2011-2 have been submitted to GenBank under the accession numbers CP116230, CP116228, and CP116229, respectively.

## Results and Discussion

After strain 18102011 was cultured overnight at 37 °C on BHI agar (with an imipenem concentration of 4 μg/mL), 0.5 mm round protruding colonies with smooth and regular edges were observed with non-fusion growth, pyocyanin production, and the absence of metallic sheens. Strain 18102011 was further identified as *P. aeruginosa* via the BD Phoenix-100 identification system and sequencing of the *oprL* gene, followed by subsequent analysis of its drug resistance spectrum (Table 1). This strain’s genome (sequence results provided in Table 2) revealed an ANI value of ≥95% with the reference strain *P. aeruginosa* POA1 (GenBank ID: NC_002516.2).

**TABLE 1.**
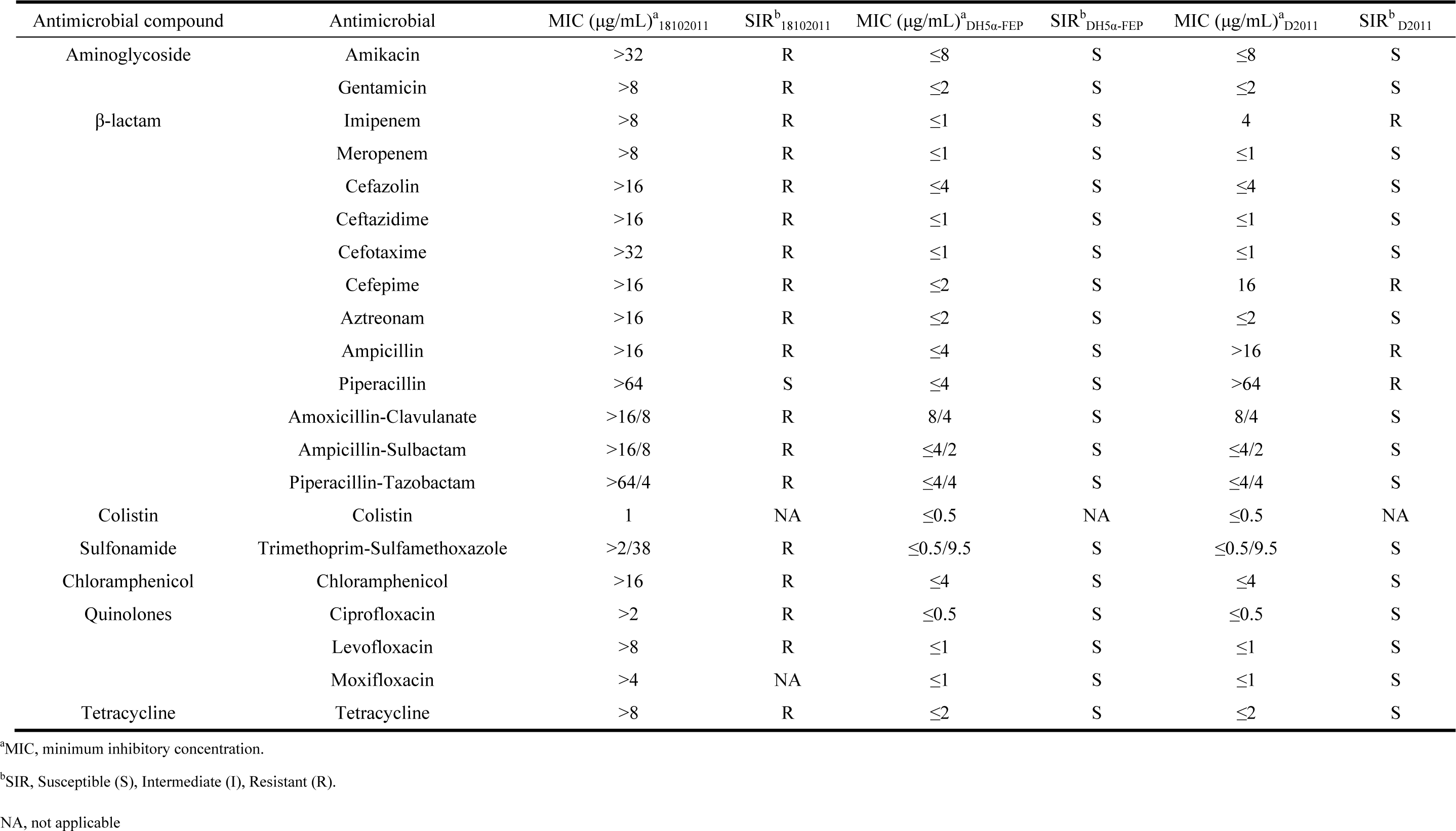
Antimicrobial susceptibility of strains 18102011, DH5α-FEP, and transconjugant D2011.

**TABLE 2.**
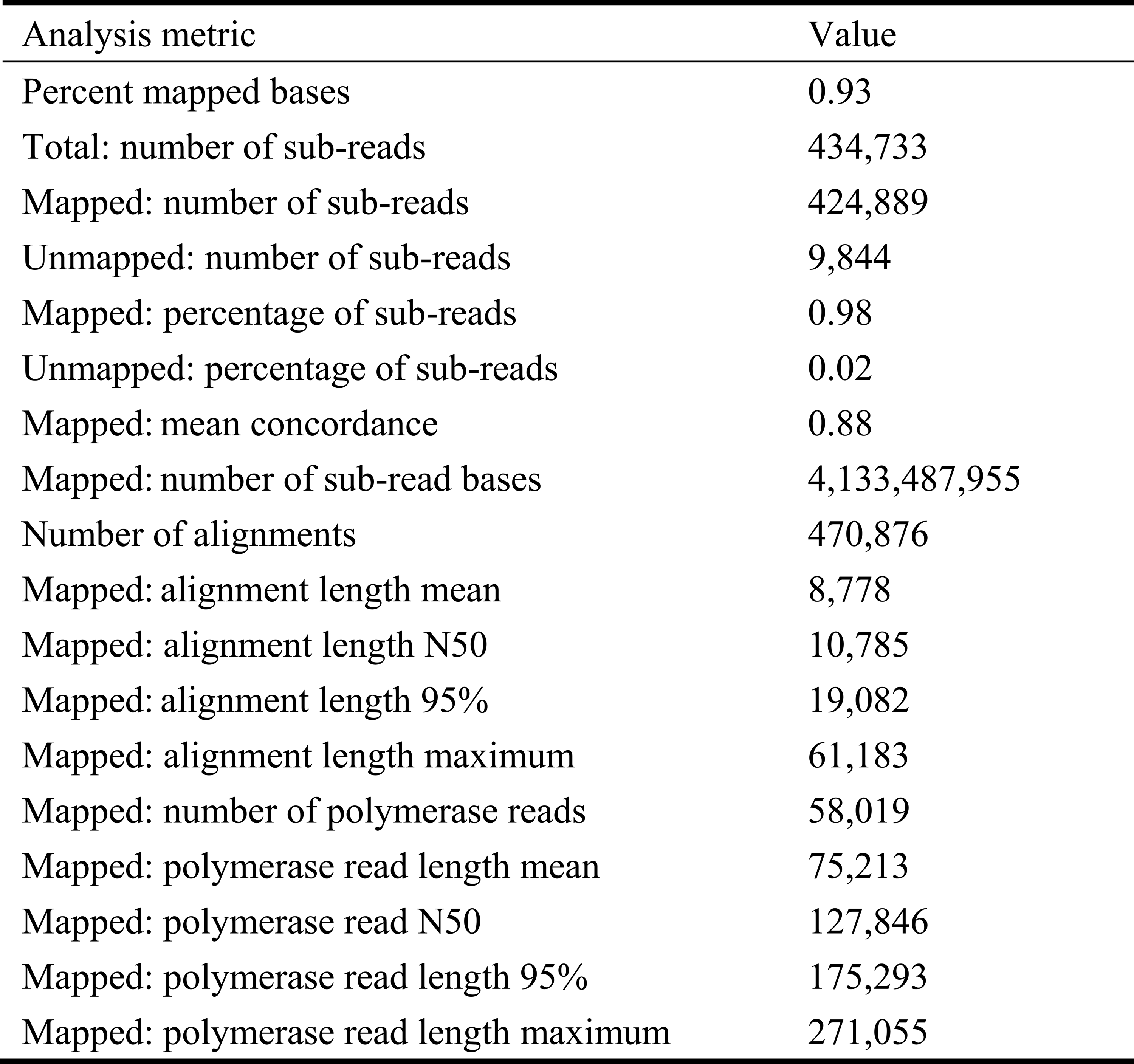
Sequencing statistics for the *P. aeruginosa* 18102011 genome.

The isolate belonged to the multi-locus sequence type ST2374 and serotype O4 based on MLST and PAst identification, respectively. The MLST database contained two total strains of ST2374 *P. aeruginosa* as of 2022, in addition to strain SX69 from a sputum origin and strain 127Gr isolated from soft tissue in Belarus in 2016. Twenty serotypes of *P. aeruginosa* are known, of which, serotype O4 is not known as a multidrug-resistant serotype ^19^. However, serotype switching from O4 to the multidrug-resistant serotype O12 has been observed under specific conditions ^20^.

Strain 18102011 carried two types of virulence genes: phospholipase C (PLC) gene (*plcH*) and Exotoxin genes (*exoS* and *exoT*). PLC is a thermolabile hemolysin that can degrade phospholipid surfactants and reduces surface tension so that alveoli do not completely collapse when air leaves them during breathing ^21,22^. The ExoS and ExoT can be obtained by the type III secretion system (T3SS) and can disrupt cytoskeletons, induces host cell rounding, disrupts intercellular tight junctions, prevents wound healing, and inhibits bacterial internalization into epithelial cells and macrophages ^23,24^. Neither of the two strongest pathogenic virulence genes-namely, the gene *exoU* encoding ExoU (T3SS effector) and *exoA* gene encoding Exotoxin A (ExoA) were not identified in this study ^25–27^, consistent with previous reports that strains do not carry both ExoS and ExoU ^28^. We speculate that strain 18102011 without ExoY (a T3SS effector) may exhibit greater cytotoxicity to epithelial cells than strains secreting active ExoY, as ExoY has been found to possibly exert a protective role at certain stages of bacterial infection, either to facilitate host colonization or to establish and/or maintain chronic infection of the host ^29^. Thus, this strain might cause cellular damage in immunocompromised patients, but it did not cause acute toxicity.

The h-CRPA 18102011 strain was resistant to all evaluated antibiotics and the MIC of IP to the strain was 4,096 μg/mL. Antibiotic resistance tests were performed with 20 antibiotics (Table 1). Aminoglycoside resistance genes (*aph(3’)-IIb*, *aac(6’)-Ib-cr*, *ant(2’’)-Ia*), β-lactam resistance genes (*bla*_OXA-396_, *bla*_PAO_, *bla*_KPC-2_, *bla*_VIM-2_, *bla*_PER-1_), a sulphonamide resistance gene (*sul1*), a chloramphenicol resistance gene (*catB7*), a quinolone resistance gene (*qnrVC6*), and a fosfomycin resistance gene *(fosA*) were identified in the h-CRPA 18102011 strain using the resfinder platform.

Genomic sequencing (summary in Table 2) revealed the presence of a 6.6 Mb chromosome of strain 18102011 (GenBank ID: CP116230) that exhibited a GC content of 66.2% and the existence of two plasmids.

The mega-plasmid pP2011-1 (GenBank ID: CP116228) carrying *bla*_VIM-2_ was 474 kb and exhibited a GC content of 56.9%. The plasmid harbored a *repA* (replication initiation) gene sharing ≥96% nucleotide identity to *repA*_IncpRBL16_. The Inc_pRBL16_ plasmid was first reported in mega-plasmid p12969-DIM (GenBank ID: KU130294) of strain *P. aeruginosa* 12969 ^30^. Including the pP2011-1 investigated in this study, only 18 sequenced Inc_pRBL16_ plasmids have been deposited in GenBank as of December 2021 (Table S1). The hosts of these plasmids were all *Pseudomonas* spp. that were isolated from environmental samples (i.e., sludge, soil, and *Origanum marjorana* samples), in addition to patient samples (e.g., from sputum, urine, or bile), primarily from China, although some instances were observed in the Netherlands and Switzerland (Table S1).

As shown in Figure 1, the backbone of pP2011-1 contained genes for partitioning (*parB2-parAB*), conjugal transfer (*cpl* and *tivF*), chemotaxis (*che*), pilus assembly (*pil*), and tellurium resistance (*ter*), in addition to *repA*_IncpRBL16_ (Figure 1). The plasmid also contained 3 novel recombination sites in addition to the common recombination site 2 for Inc_pRBL16_ family plasmids ^31^. IS*Pa75* (IS*66* family) and *fosA* formed the accessory module 1 region that was inserted between two hypothetical proteins of the tellurium resistance region. In addition, *dnaK* and *yaeT*, along with several phage integrase genes and stress protein genes formed the accessory module 2 region that was inserted into the region of the catalytic subunit of Pol V (UmuC), which is a major recombination site of Inc_pRBL16_ family plasmids. The accessory module 3 region comprised heavy-metal efflux protein genes (*merE*, *merD*, *merA*, *merP*, *merT*, and *merR*), Tn*6001* (Tn*3* family), two copies of IS*CR1* (IS*91* family), *bla*_PER-1_, *qnrVC6*, and a novel type I integron In2057 carrying *ant(2’’)-IIa*, *bla*_VIM-2_ and *aac(6’)-Ib-cr*, which was inserted between the hypothetical protein genes that were 708 kb and 1,257 kb in length. Lastly, the accessory module 4 region included IS*Pa141* (IS*30* family), IS*Pa61* (IS*L3* family), IS*Pst3* (IS*21* family), IS*Pa60* (IS*As1* family), Tn*4662a* (Tn*3* family), and Tn*5046.1* (Tn*3* family), which was inserted between the hypothetical protein and phage protein genes.

**Figure 1.**
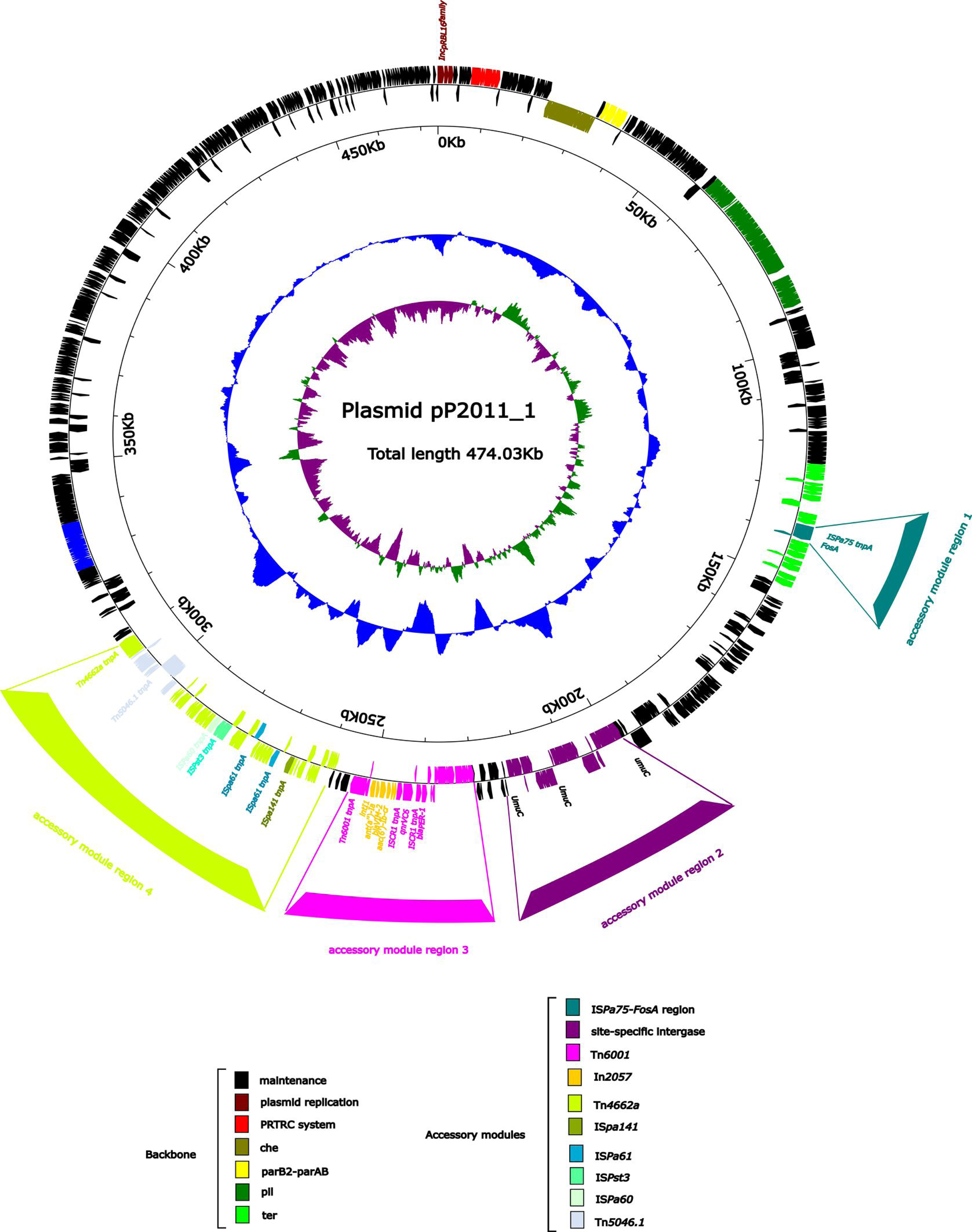
Annotation of the plasmid pP2011-1. Circles show (from outside to inside): (1) genome functional annotation. The backbone region is black, the plasmid replicon is brown, the PRTRC system is red, the *che* region is dark green, the parB2-parAB region is yellow, the *pil* region is lighter green, and the *ter* region is bright green. IS*Pa75* and the *fosA* genes form the accessory module 1 region (dark cyan), the *dnaK* and, *yaeT* genes, along with several phage integrases genes and stress protein genes, form the accessory module 2 region (purple) that was inserted into UmuC region (purple). The accessory module 3 region (pink) comprised Tn*6011* (pink) and In2057 (orange). The accessory module 4 region (green-yellow) comprise IS*pa141* (olive), IS*pa61* (dark blue), IS*pst3* (turquoise), IS*Pa60* (honeydew), Tn*4662a* (green-yellow) and Tn*5046.1* (alice blue); (2) GC skewcalculated as [(G-C)/(G+C)]; and (3) GC content.

As shown in Figure 2, plasmid pP2011-2 (GenBank ID: CP116229) carrying *bla*_KPC-2_ was 40 kb in length and exhibited a GC content of 58.1%. The plasmid harbored a *repA* gene sharing ≥96% nucleotide identity to *repA*_Incp6_. The backbone of the *repA*_Incp6_ family plasmid included *repA*_Incp6_, genes for partitioning (*parABC*) and a mobilization region (*mobABCDE*), in addition to two accessory modules. The accessory module 1 region comprised Tn*5563a* (Tn*3* family) and IS*Pa19*, while accessory module 2 included IS*Ec33*-Tn*3*-IS*Apu2*-IS*Apu2*-*orf7*-IS*Kpn27*-*bla*_KPC-2_-ΔIS*Kpn6-KorC-orf6-klcA-*Δ*repB*. The IncP6 plasmid was first identified in pRms149 (GenBank ID: GCA_019400855.1) of the strain *P. aeruginosa* JX05 in 2005 ^32^. Including pP2011-2, only 27 sequenced Incp6 plasmids carrying Tn*3*-ΔIS*Kpn6*-*bla*_KPC-2_-IS*Kpn27* have been deposited as of December 2021 in GenBank (Table S2). The hosts of these plasmids primarily were Enterobacteriaceae, but also included some *P. aeruginosa* and *Aeromonas* strains (Table S2). Further, the hosts primarily derived from China, but also included some examples from Japan, Germany, the United States, Argentina, Croatia, and Vietnam, among other countries. The hosts were isolated from sewage and patient samples (e.g., from blood, urine, and bile) (Table S2). The rapid spread of *bla*_KPC_ genes has been associated with their localization with Tn*4401*, a Tn*3*-based transposon capable of high-frequency transposition that itself is present on diverse enterobacterial plasmids varying in size, nature, and structure. The Tn*3*-IS*Kpn27*-*bla*_KPC_-ΔIS*Kpn6* structure has been commonly identified as the *bla*_KPC_ platform in China, but not Tn*4401* ^33^. To generate the above core module, an intact IS*Kpn27* would have been inserted into Tn*3*, forming Tn*3*-IS*Kpn27* that was then linked with *bla*_KPC_ and ΔIS*Kpn6* ^34^.

**Figure 2.**
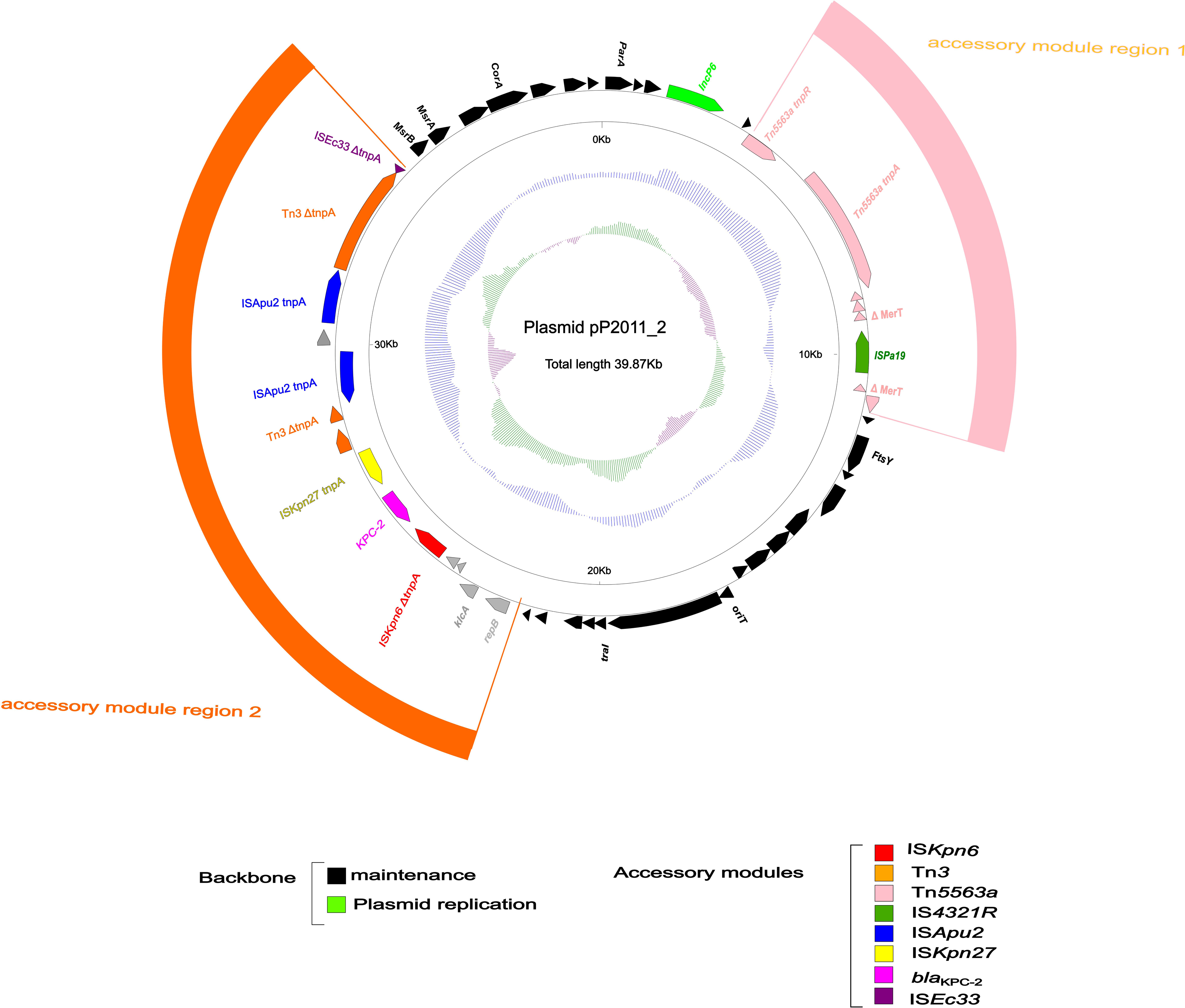
Annotation of plasmid pP2011-2. Circles show (from outside to inside): (1) genome functional annotation. The backbone region is black, in which the plasmid replicon is green. Tn*5563a* (pink) and IS*Pa19* (sea-green) comprise accessory module 1 region (pink), while the module 2 region (orange) comprise IS*Kpn6* (red), *bla*_KPC-2_ (magenta), IS*Kpn27* (yellow), Tn*3* (orange), IS*Apu2* (blue), and IS*Ec33* (purple); (2) GC skew calculated as [(G-C)/(G+C)]; and (3) GC content.

Tn*3*-ΔIS*Kpn6*-*bla*_KPC-2_-IS*Kpn27* has previously been shown in association with *korC*-*orf6*-*klcA*-Δ*repB* to form the major genetic structure (Tn*3*-ΔIS*Kpn6*-*bla*_KPC-2_-IS*Kpn27-korC*-*orf6*-*klcA*-Δ*repB*) inserted into the Tn*1722* site of pKP048 (GenBank ID: NC_014312.1) ^34^. The structure has also been excised from Tn*1722* and reorganized upstream of ΔIS*Ec33*, forming the ΔIS*Ec33* associated-element ΔIS*Ec33-Tn3-ISApu1-orf7-ISApu2-ISKpn27-*Δ*bla*_TEM-1_-*bla*_KPC-2_-ΔIS*Kpn6-KorC-orf6- klcA-*Δ*repB* in p10265-KPC (GenBank ID: KU578314.1) ^34^. pP2011-2 and p10265-KPC all harbored one copy of IS*Pa19* and a 7.9 kb Tn*5563a* transposon, with IS*Pa19*_p10265-KPC_ inserted downstream of Tn*5563*_p10265-KPC_ and IS*Pa19*_pP2011-2_ inserted inside Tn*5563*_pP2011-2_, resulting in truncation of its *merT* (Tn*5563*_pP2011-2_), and thus Tn*5563*_pP2011-2_ structure formation occurring after Tn*5563*_p10265-KPC_. Further, pP2011-1 did not contain Δ*bla*_TEM-1_, which may have been excised during horizontal transfer of the plasmid p10265-KPC.

Conjugation experiments revealed that the plasmids pP2011-1 and pP2011-2 of *P. aeruginosa* could be transferred into *E. coli*, with an associated conjugation transfer efficiency of 10^-6^. The *E. coli* transconjugant carried *bla*_KPC-2_ and *bla*_VIM-2_ from the donor, and the MIC value of IP to the *E. coli* transconjugant was 4,096 μg/mL, which was the same as the MIC value for the donor. Co-harboring of different carbapenemase gene types in *K. pneumoniae* has been reported with increasing frequency ^35,36^, while, *P. aeruginosa* co-harboring different types of carbapenemase genes has also been identified in many countries ^37,38^. In contrast to previous studies, the strain 18102011 evaluated in this study was derived from a bile sample, representing the first instance of an isolate from bile co-harboring different types of carbapenemase genes.

## Conclusions

Here, strain ST2374 corresponding to an h-CRPA isolate from an ICU patient bile sample was characterized and identified to carry *bla*_KPC-2_ and *bla*_VIM-2_. The strain harbored a novel type 1 integron containing *bla*_VIM-2_ through the Inc_pRBL16_ mega-plasmid pP2011-1 that contained 3 novel recombination sites. Concomitantly, *bla*_KPC-2_ was identified in a IS*Ec33*-Tn*3*-IS*Apu2*-IS*Apu2*-*orf7*-IS*Kpn27*-*bla*_KPC-2_-ΔIS*Kpn6-KorC-orf6-klcA-*Δ*repB* structure in the IncP6 type plasmid pP2011-2. The presence of different types of carbapenemase genes represent challenges for treating infections by their host bacteria. Hence, the co-harboring of different types of carbapenemase genes in bacteria should be closely monitored and evaluated epidemiologically across the world.

## Author contributions

All strains were provided by China-Japan Union Hospital, Jilin University. XJG, LWZ, YLY and PC conceived, directed, and carried out the study. JYG, QLL, GJL, YS, XJ, JJ, SWS, and BWJ prepared samples for sequence analysis. JYG, LZ, and ZXW acquired samples and analyzed the data. LZ and ZXW conducted this manuscript. All authors have read and approved the final manuscript.

## Acknowledgments

We are grateful to the members of the China-Japan Union Hospital, Jilin University.

